# Pangenome reconstruction of *Lactobacillaceae* metabolism predicts species-specific metabolic traits

**DOI:** 10.1101/2023.09.18.558222

**Authors:** Omid Ardalani, Patrick Phaneuf, Omkar S. Mohite, Lars K. Nielsen, Bernhard O. Palsson

## Abstract

Strains across the *Lactobacillaceae* family form the basis for a trillion-dollar industry. Our understanding of the genomic basis for their key traits is fragmented, however, including the metabolism that is foundational to their industrial uses. Pangenome analysis of publicly available *Lactobacillaceae* genomes allowed us to generate genome-scale metabolic network reconstructions for 26 species of industrial importance. Their manual curation led to more than 75,000 gene-protein-reaction associations that were deployed to generate 2,446 genome-scale metabolic models. Cross-referencing genomes and known metabolic traits allowed for manual metabolic network curation and validation of the metabolic models. As a result, we provide the first pangenomic basis for metabolism in the *Lactobacillaceae* family and a collection of predictive computational metabolic models that enable a variety of practical uses.

## Introduction

*Lactobacillaceae* are an essential family of highly diverse lactic acid bacteria. It comprises a large number of species that populate a variety of habitats *(1)*. Due to numerous applications in food and pharmaceutical industries, *Lactobacillaceae*-dependent products have a trillion-dollar market size, including dairy, wine, probiotics, and numerous satellite industries (2–4) (see Supplementary Table 2 for details), indicating their importance in microbial biotechnology and related industries.

Cost-effective DNA sequencing has led to a steadily increasing number of *Lactobacillaceae* genomes deposited in the NCBI database (5). The availability of these genomes has enabled pangenomic studies (1,6–8). Metabolism is foundational to the industrial uses of *Lactobacillaceae* and the generation of predictive genome-scale metabolic models (GEMs) constitutes a significant advancement in bioprocess engineering (9). GEMs are based on annotated sequences and use algorithms to forecast cellular behavior and metabolic fluxes under conditions of interest. GEMs have been shown to predict optimal growth conditions and gene manipulation targets to increase product yield by integrating complex metabolic pathways. The availability of a large number of *Lactobacillaceae* genome sequences enables pangenome-based metabolic reconstruction to create a comprehensive and predictive set of GEMs.

In this study, 2,446 GEMs were generated across the *Lactobacillaceae* family. This set of models (called a PanGEM) enables the identification of conserved and variable metabolic traits across different family members. The individual GEMs can be used to develop strategies for strain improvement and bioprocess optimization. Moreover, predictive GEMs have a wide range of applications, from understanding the metabolic basis for probiotic properties to producing value-added compounds for the food and pharmaceutical industries. Overall, constructing the PanGEM is important to fully harness *Lactobacillaceae*’s biotechnological potential, and it constitutes a new quantitative representation of the genomic basis for this large industry.

## Results

### Metabolic network reconstruction and genome-scale models for a family of bacteria

First, a metabolic pan-reactome was constructed for use as a template for species-specific metabolic reconstructions. The pan-reactome was reconstructed from 49 high-quality reference genomes, representing 33 species from 9 genera (Supplementary File 1). The reactome (Supplementary File 2) contained 75,299 gene-protein-reaction associations (GPRs), 1,873 reactions, 28,280 genes, and 1,659 metabolites (Fig.1a, stage 1). Reactions were distributed across 231 cellular subsystems. Out of 1,873 reactions, 1,516 were gene-associated reactions, whereas the remaining belonged to other types, including exchange, orphan, gap (see Supplementary Figure 8 for a detailed gap analysis), demand, and sink reactions.

**Figure 1.**
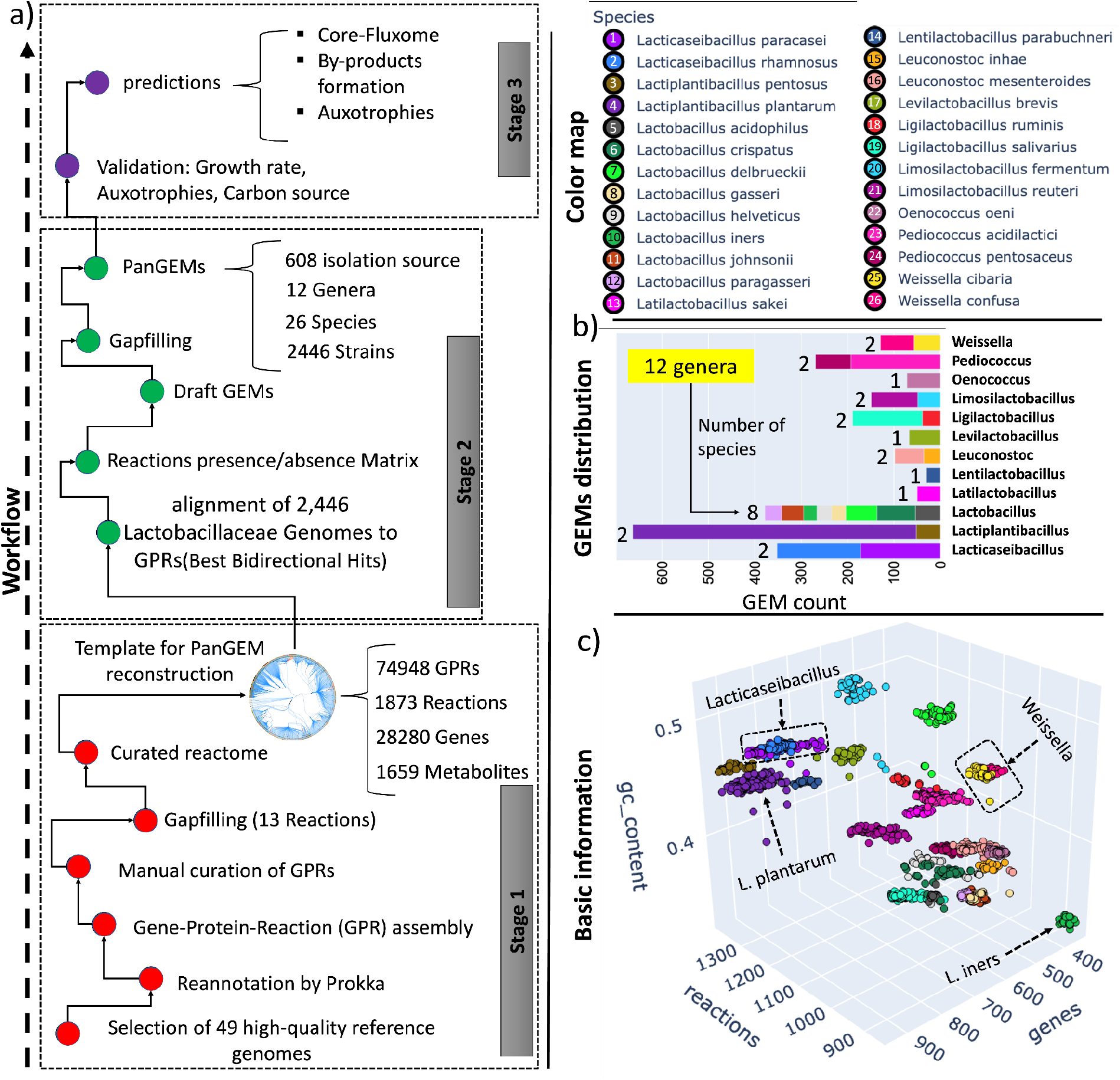
Pangenome-scale metabolic reconstructions for Lactobacillaceae. **a)** the overall three-stage workflow used in this study. Stage 1: Reactome reconstruction. To formulate a draft pan-metabolic network reconstruction, 49 Lactobacillaceae reference genomes were selected and annotated by Prokka. The draft reactome was manually curated to develop ‘gold-standard’ multi-allelic gene-protein-reaction associations (GPRs), covering 1,832 reactions and 28,280 metabolic alleles. Stage 2: Building strain-specific metabolic reconstructions on a family-wide basis. 2,446 Lactobacillaceae genomes from NCBI were aligned with BLAST against the GPRs to identify the corresponding reactions based on sequence similarity. A biomass objective function was added to all draft reconstructions to identify and fill gaps in the metabolic reconstruction. At this step, all GEMs were able to produce biomass and were ready for further analysis. Stage 3: Validation and analysis of strain-specific GEMs. Additional validation through iterative refinement was performed to achieve precise and accurate metabolic phenotype predictions. After GEMs passed quality control, analysis was performed to identify common metabolic capabilities and differences across individual Lactobacillaceae strains. **b)** GEM distribution across the Lactobacillaceae. GEMs are classified based on their genus (Y-axis). Different species within each genus are annotated according to the color map; bar plots are annotated with the number of species within each genus. PanGEM contains 12 genera, including Lactobacillus, Ligilactobacillus, Limosilactobacillus, Lacticaseibacillus, Oenococcus, Leuconostoc, Levilactobacillus, Pediococcus, Lactiplantibacillus, Latilactobacillus, Weissella, Lentilactobacillus. **c)** Depicts the basic information about the metabolic reconstructions in PanGEM. Color map annotates species, and the three axes depict the number of reactions, number of genes for each reconstruction, and GC content of the corresponding genome. These three parameters give distinct clusters for the stains of each species.

Second, metabolic models for the *Lactobacillaceae* family were constructed. The GPRs were matched against the ORFs in every qualified genome, resulting in 2,446 strain-specific GEMs from 26 species and 12 genera obtained from 608 distinct isolation sources (6) (Supplementary Figure 1). *L. plantarum* had the highest number of GEMs (611), while *L. iners* and *L. parabuchneri* had the lowest (30 each). The genus *Lactiplantibacillus* had the most GEMs (663), while *Lentilactobacillus* had the least (30) (see Fig.1b). In the *Lactobacillaceae* PanGEM, the genus *Lactobacillus* had the highest diversity with eight species, while *Oenococcus, Levilactobacillus, Lentilactobacillus*, and *Latilactobacillus* had only one species each, resulting in the lowest diversity within the group.

The PanGEM covers a wide range of metabolic diversity across 26 *Lactobacillaceae* species, with multiple GEMs for each species. The number of reactions per genome ranged from 859 to 1,358, representing 354 to 944 genes, respectively. On average, *L. iners* had the lowest number of genes and reactions, while *L. pentosus* had the highest (see Supplementary Figure 2). Each species can be distinguished from other species based on three basic properties (number of genes, number of reactions, and GC-content of genomes) (Fig.1c). Also, PCA analysis revealed distinct metabolic profiles across the various species of *Lactobacillaceae*, as evidenced by the observed clustering patterns which indicate adaptations to various ecological niches (see Supplementary Figure 3 for more information). The PanGEM is the first of its kind, a collection of metabolic models across an entire family.

### Validation of PanGEM

Once formulated, GEMs were validated against experimentally determined metabolic and physiological traits (10). Growth predictions on 21 carbon sources for three strains of *Limosilactibacillus* (sakei DSM 20017, sakei LS25, sakei 23k) were compared to experimental results available in the literature (11) (Fig.2a). Out of 63 simulations, 52 (83%) were accurate, while two were false negatives, and nine were false positives. Both predictions and experimental results were obtained on chemically defined media (CDM). (see CDM formulation and constraints in Supplementary Table 1).

**Figure 2.**
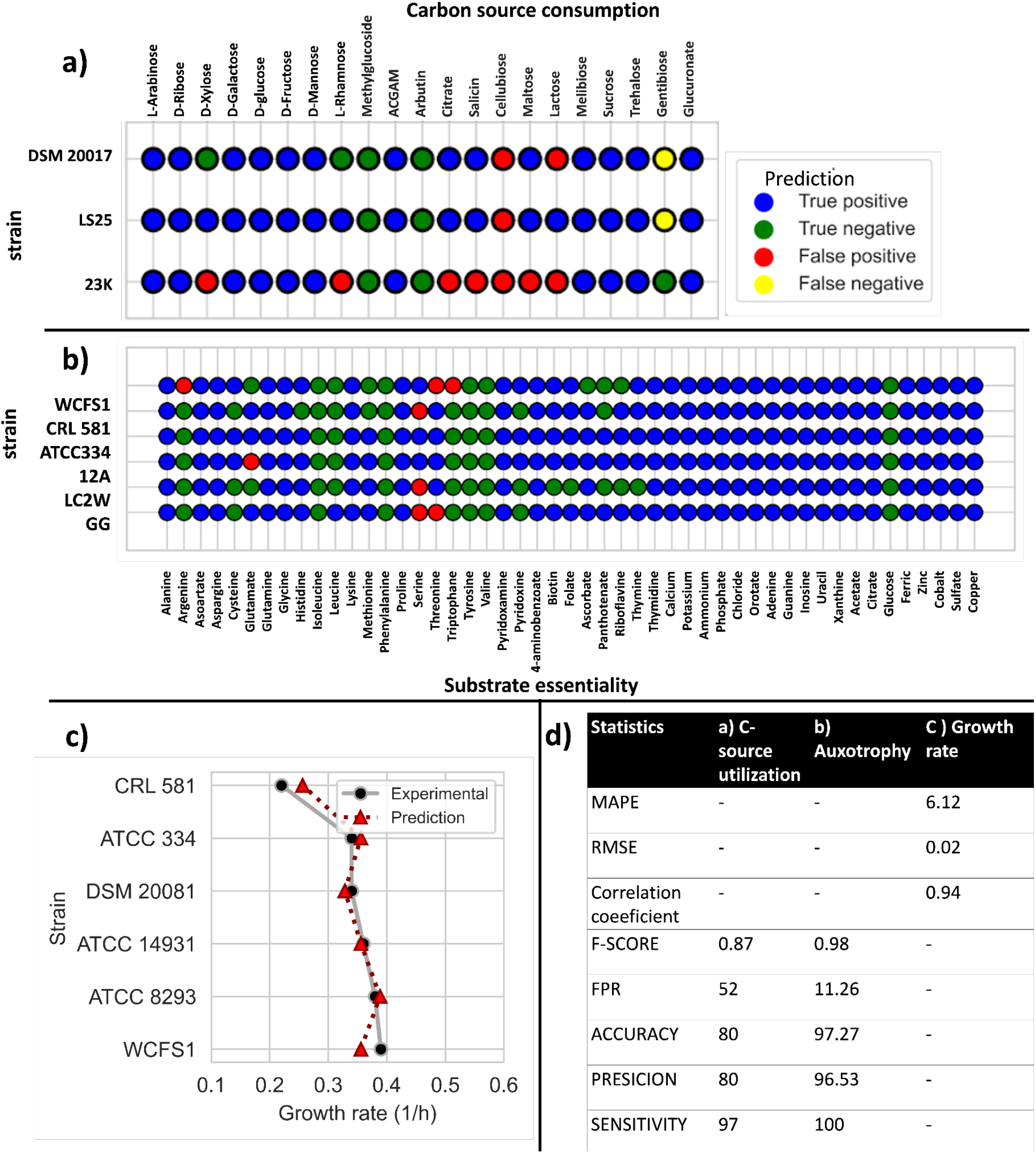
PanGEM validation. **a)** Carbon source utilization. Carbon source utilization validation for three strains within PanGEM (L. sakei 23K, L. sakei DSM 20017, L. sakei LS25) by comparing experimental and predicted results. The capability to grow on 21 carbon sources (X-axis) for three strains (Y-axis) is shown; the color map annotates False/True/Positive/Negative predictions. Statistical analysis based on a confusion matrix showed an F-score of 0.98, a false positive rate (FPR) of 11.26%, a precision of 96.53%, an accuracy of 97.27%, and a sensitivity of 100% **b)** Prediction of auxotrophies. Validation of PanGEM by comparing experimental and predicted results for chemically defined medium (CDM) component essentiality for six strains (L. plantarum WCSF1, L. delbrueckii CRL581, L. paracasei ATCC334, L. paracasei LC2W, L. rhamnosus GG (Y axis)). A single-component omission analysis of 49 components of Lactobacillaceaei-specific CDM was simulated using FBA, and results were compared with experimental data. Orange dots represent false positive predictions. Blue and green dots represent true negative and true positive, respectively. No false negative prediction was observed, with an F-score of 0.87, a false positive rate (FPR) of 52%, a precision of 80%, an accuracy of 80%, and a sensitivity of 97%. **c**) A comparison of simulated growth rate using FBA on CDM (dashed line-red triangle) with experimental data on the same condition (solid line-black dots) for six strains within PanGEM. (Abbreviations: CRL 581; L. delbrueckii CRL 581, ATCC 8293; L. mesenteroides ATCC 8293, DSM 20017; L.sakei DSM 20081, DSM 20017; L.sakei DSM 20081, LS25; L.sakei LS25, 23K; L.sakei 23K, WCFS1; L. plantarum WCFS1, ATCC334; L. paracasei ATCC334, 12A; L. Paracasei 12A, LC2W; L. Paracasei LC2W, GG; L. Rhamnosus GG, ATCC 14931; Limosilactobacillus fermentum ATCC 14931). **d)** Statistics. Statistical analysis of GEM validation results is summarized in the table. For quantitative predictions (growth rates), MAPE, RMSE, and correlation coefficient were calculated. For qualitative predictions (auxotrophy and C-source utilization), F-score, FPR, precision, accuracy, and sensitivity were calculated.

For further qualitative phenotypic validation of the PanGEM, single omission analysis of 49 compounds in a chemically defined medium (CDM) was simulated by flux-balance analysis (FBA). The computational results were compared to experimental data (12–16) (Fig.2a2). Experimental data were available for six strains within PanGEM, representing four different species and three unique genera. FBA was performed to predict the single omission of 49 compounds from CDM, one-by-one across six strains. Among 294 simulations, only 8 (2.7%) failed to predict the correct phenotype (False positives) (Fig.2b).

Growth rate computations showed that the models were capable of quantitative prediction of growth rates that were in agreement with the experimental reports (13,17–19) (Fig.2c). Despite the limited growth rate data available in the literature from CDM, validation was performed on six GEMs, and predictions were satisfactory with a mean absolute percentage error (MAPE) of 6.62, root mean squared error (RMSE) of 0.02 and correlation coefficient of 0.93. These metrics illustrate the prediction potential of PanGEM models (Fig.2c).

PanGEM could thus be validated against data reported in the literature. A fundamental reason for panGEM’s predictive power is the relative simplicity of *Lactobacillaceae* metabolism and that the GPRs reflect its genomic basis well. The validated PanGEM can be used for the analysis of more detailed metabolic traits.

### Classifying strain-specific reactomes in PanGEM

Strain-specific GEMs can be clustered based on the reactions they contain (Fig.3a). Such clustering highlights metabolic differences between species. The cluster map (Fig.3a) shows three clusters of reactions; core reactions (common to all strains), accessory reactions (found in many strains), and rare reactions (found in few or even in single strains, then called unique). All strains shared 183 common reactions (Fig.3b), forming the *Lactobacillaceae* core-reactome. There were 1130 accessory reactions, and they are differentially distributed across the 26 strains (Fig 3a). Metabolic conservation in each species was assessed by performing a reaction frequency analysis (Fig.3d). Results indicate that *L. plantarum* had the most metabolically diverse strains, while *L. acidophilus* was the most metabolically conserved species (Supplementary Note 1).

**Figure 3.**
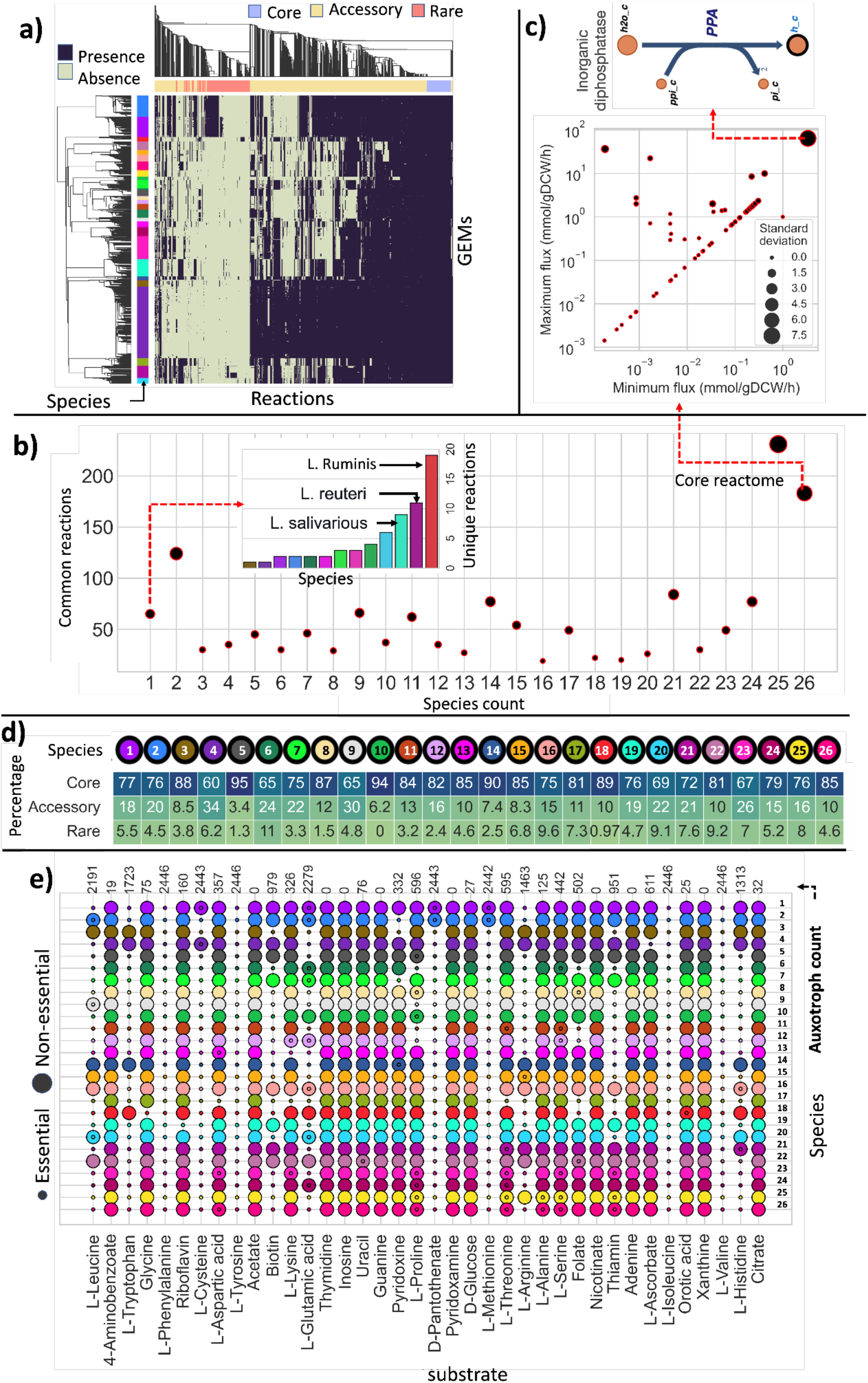
Characteristics of the Lactobacillaceae Reactome. **a)** Reaction Presence-absence cluster map shows reaction presence and absence calls in each strain (represented by a row) in each species (color-coded as a group of rows). Strain-specific metabolic network reconstructions and reactions are represented by rows and columns, respectively. Row color refers to species; column color represents core, accessory, and rare reactome. **b)** Shared reactions distribution across the 26 species. Scatter plot depicts the number of common reactions among different species., The inset bar chart represents the unique reaction count per species. Bars are annotated based on the species color map (see Species color map in Fig.1). **c)** Core-fluxome differential analysis defines metabolic flexibility. Scatter plot zooms in on common reactions across all species (core-reactome). Dots represent reactions, predicted minimum (X-axis) and maximum (Y-axis) fluxes represent strains with lowest and highest flux(mmol/gDCW/h) for each reaction. Dot size shows standard deviation of fluxes through each reaction across all strains. Demands, sinks, and exchanges, reactions were excluded from core-fluxome **d)** Intra-species reactions frequency analysis. Heatmap depicts species-specific core, accessory, and rare reactome, color dot represents species (see fig.1 species color map), and values within the heatmap are the percentage of intra-species core, accessory, and rare reactome. **e)** PanGEM prediction of species auxotrophies in CDM.

65 unique reactions were found within 13 species (Fig.3b, bar chart). Among all species, *L. ruminis*, a niche-specialized species, had the highest unique reaction count of 19. Next was *L. reuteri* with 11 unique reactions, followed by *L. salivarius*, with nine unique reactions (Supplementary Figure 4).

**Figure 4.**
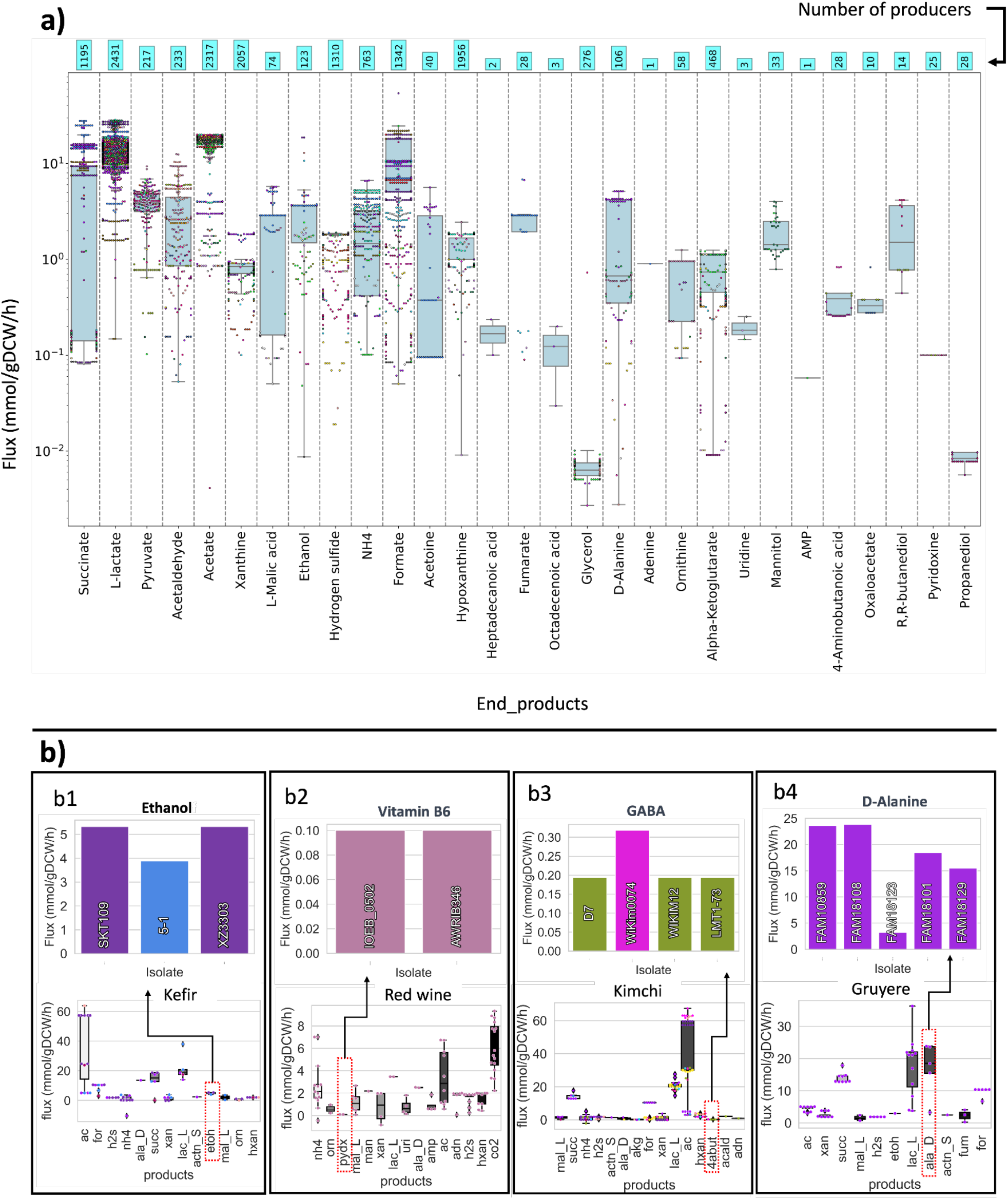
PanGEM prediction of by-product formation phenotypes. **a)** Isolation source-specific by-product profiling. Box plots depict the predicted production profile of strains from four different isolation sources, and color dots represent different species (see Fig.1 species color map). Bar charts zoom in on isolates responsible for the production of key metabolites. Bars are colored based on species color map. b) Prediction of by-product formation across PanGEM models. The box plot depicts the capability of different species in product formation on CDM. Colored dots represent species and are coded (see Fig.1 species color map). b1) Tibet kefir isolates metabolite production profile, b2) Red wine isolates metabolite production profile, b3) Kimchi isolates metabolite production profile and b4) Gruyere (a Swiss cheese) isolates metabolite production profile. An important by-product for each isolation source is selected by the dashed line to depict producer strains and their production rates regarding the selected by-product (bar plots pointed by dashed arrows).

### Characterization of allowable flux states using PanGEM

FBA analysis was performed for all 2,446 strain models in PanGEM to better understand Lactobacillaceae core metabolism. Among 183 common core reactions, a subset of 142 reactions belonging to 30 different metabolic subsystems was found to represent common active reactions, which means they could consistently carry metabolic flux in all strains. Therefore, they were called the *Lactobacillaceae* core-fluxome (Supplementary File 3). The remaining 41 reactions from the core-reactome that were excluded from the core-fluxome were not carrying any flux.

Among the 30 subsystems of the core-fluxome, arginine biosynthesis, cysteine, methionine metabolism, fatty acid biosynthesis, and purine metabolism have the highest fluxes. Other subsystems within the core-fluxome were mostly involved in biomass production, such as pyrimidine metabolism, peptidoglycan biosynthesis, aminosugar metabolism, and glycerophospholipid biosynthesis. Nicotinate and nicotine metabolism and pantothenate and CoA biosynthesis were the only subsystems related to cofactor biosynthesis within the core-fluxome.

FBA was performed to assess the variability of fluxes through each reaction across all strains in the core-fluxome. Results suggest that the core-fluxome is consistent within all *Lactobacillaceae* and carries a very similar metabolic flux for all strains. To find the reactions with the most variable flux across the PanGEM, the standard deviation of minimum and maximum fluxes was calculated (Fig.3c). Among the core-fluxome, inorganic diphosphatase (PPA) had the most variable fluxes ranging from 3.3 to 63.2 mmol/gDCW/h, with an average value of 15.4 mmol/gDCW/h and standard deviation of 7.6 mmol/gDCW/h. Further analysis revealed a correlation between the growth rates of strains and the flux through the PPA reaction (Supplementary figure 5).

FBA-predicted CDM media component essentiality for each GEM was used to reveal metabolic similarities and differences in *Lactobacillaceae* (Fig.3e). *L. plantarum* had the lowest auxotrophy count. *Pediococcus acidilactici* and *cibaria* had the highest number; *L. sakei* and *L. paragasseri* had the most consistent auxotrophies. Isoleucine, valine, phenylalanine, and tyrosine were globally essential in all 2,446 models in PanGEM (Supplementary Note 2).

The PanGEM enabled the assessment of the composition of the reactome and the range of allowable flux states. The most interesting finding about the reactome composition is the prevalence of species-specific unique reactions. In terms of the core fluxome, reactions exhibited consistent flux across the panGEM, while certain reactions displayed varying flux values within each individual GEM when compared to the others, such as PPA.

### Biotechnological potential of *Lactobacillaceae* can be discovered using PanGEM

Validated and characterized PanGEM models can be used for a variety of applications (20,21). Metabolic by-product secretion is a defining metabolic trait of *Lactobacillaceae* and determines their uses in food production. FBA analysis was performed to predict potential metabolite production across *Lactobacillaceae* (Fig.4). As expected, PanGEM models showed high production rates for lactate and acetate (Fig.4a). Metabolite connectivity analysis across all metabolic networks of PanGEM showed that the diversity of *Lactobacillaceae* end-products had a strong correlation with connectivity of pyruvate and glutamate (Supplementary Figure 6c).

PanGEM predicted the production of several flavor compounds, such as acetoin, acetaldehyde, pyruvate, succinate, D-alanine, and ethanol, as well as the neurotransmitter GABA. *L. paracasei* strains were predicted to be major D-Alanine producers. Ethanol producers were mostly plant-based *L. pentosus* and *L. brevis*, dairy-based *P. acidilactici*, and commensal *L. paracasei*. Succinate producers include commensal *L. rhamnosus, P. acidilactici*, and *L. paracasei* isolated from unknown sources. Gamma-aminobutyric acid (GABA) production was predicted in two species, commensal *P. acidilactici* and commensal *L. brevis* (Supplementary Figure 6a). PanGEM thus predicts the metabolic by-product formation on a species-specific basis.

To understand the biotechnological capabilities of product-specific isolates and how they contribute to desirable organoleptic properties of the final product, FBA was performed on GEMs belonging to isolates from four industrial products, including Kimchi, Kefir, Gruyere, and red wine. FBA predicted different fermentation profiles for each product. Tibet kefir includes 14 isolates from *L. paracasei, L. mesenteroides, L. plantarum, P. pentosaceus, and L. rhamnosus* that are potentially capable of producing 13 metabolites on CDM (Fig.4b1). Among these metabolites, ethanol, succinate, acetoin, and lactic acid are responsible for desirable flavors in this specific product and have been the subject of several studies. Among all kefir isolates, two *L. plantarum* and one *L. rhamnosus* isolates are responsible for ethanol production (Fig.4b1).

Red wine isolates of *O. oen*i exhibited a broader range of metabolite production. Notably, among the 15 metabolites that could potentially be produced, pyridoxine and ornithine are important in the wine industry. (Fig.4.b2).

Kimchi includes 52 isolates from ten species, including *W. cibaria, P. pentosaceus, L. mesenteroides, L. sakei, L. plantarum, L. brevis, L. fermentum, P. acidilactici, L. paracasei, and L. rhamnosus*. FBA predicted that these strains produce 15 different metabolites, among which formate, acetaldehyde, acetoin, succinate, and GABA play a crucial role in the quality of the final product. FBA showed GABA producer isolates were three *L. brevis* and one *L. sakei* isolate; *L. sakei* showed a higher GABA production rate than *L. brevis* isolates (Fig.4b3).

The final prediction was performed on 11 Gruyere isolates, all from *L. paracasei*. FBA predicted the production of 11 different metabolites; formate, acetoin, and succinate are considered important product-specific metabolites. Also, interestingly, a high production rate for D-alanine was predicted. Since Gruyere is known for its sweet taste (22), and D-alanine is a known amino acid-based sweetener (23), there might be a link between this phenotype and Gruyere’s organoleptic properties. FBA predicted that six isolates were D-alanine producers (Fig.4b4).

The pangenome metabolic reconstructions and the metabolic traits that they predict for 2,446 strains were thus highly consistent with the characteristics of the genera and species to which they belong. PanGEM thus provides a global atlas of the genetic basis for metabolic traits in the *Lactobacillaceae* family. One can select an isolation source, identify strains specifically from that source, and conduct FBA to construct an end-product profile. This profile can then be a valuable guide for directing subsequent experimental assessments.

## Discussion

*Lactobacillaceae* is widely used in pharmaceutical, fermented, and beverage industries (Supplementary Table 2). As producers of various compounds, such as lactic acid, acetic acid, succinic acid, diacetyl, acetoin, GABA, sorbitol, mannitol, butanediol, propanediol, and the vitamin B family, lactic acid bacteria have a strong potential for biotransformation. Despite their enormous industrial potential, prominent market share, and the availability of large amounts of omics data (24–27), we lack computational models that link genotypes to desirable industrial phenotypes. This study remedies this shortcoming by using pangenome analysis and metabolic reconstruction to characterize the metabolic potential of the *Lactobacillaceae* by generating 2,446 high-quality GEMs across 26 species obtained from 608 isolation sources.

Reactome analysis demonstrated that the core reactome comprises only 8% of the total reactome. In comparison, the accessory and rare reactomes account for 73% and 19%, respectively, indicating significant diversity in metabolic capabilities among *Lactobacillaceae*. Of the 65 unique reactions identified in the reactome, *L. ruminis, L. reuteri*, and *L. salivarius* have the highest number. These species commonly reside in the human gut and oral cavity, suggesting that acquiring novel metabolic capabilities may be crucial for bacterial adaptation and survival in complex environments. *Lactobacillus ruminis* demonstrates the highest count of unique reactions, comprising a total of 19. Certain unique reactions, such as TEICEXP, TEICEXP2, and TEICEXP3 related to teichoic acid export, were only found in *Pediococcus acidilactici*, which may be a distinct trait of this species. Previous studies have indicated that the thick cell wall of *P. acidilactic*i plays a crucial role in heavy metal accumulation in the gut (28). *P. acidilactici and L. brevis* were also responsible for approximately 70% of microbial spoilage in beer due to hop resistance linked to the cell wall and teichoic acid structure (29). Additionally, choline trimethylamine-lyase was found only in *L. plantarum* strains. This enzyme cleaves choline to produce trimethylamine (TMA) and acetaldehyde. TMA causes several disease-associated microbial metabolites *(30)*. Therefore, PanGEM could screen for disease-associated microbial metabolites such as TMA and their producers to avoid or reduce their usage in food industries.

Analysis of the core-fluxome shows the possible activity states of core metabolism. Its properties revealed a consistent flux range across *Lactobacillaceae*. However, inorganic pyrophosphatase (PPA) showed the most flux variation across different strains. PPA is a crucial enzyme for cellular homeostasis since many essential anabolic pathways produce pyrophosphate (PPi). PPi hydrolysis maintains anabolic pathways by recycling orthophosphate (31). We found a significant positive correlation between the predicted flux of PPA and the predicted growth rates of the strains, highlighting the significance of PPA in the metabolism of *Lactobacillaceae*.

FBA predicted global auxotrophy, including isoleucine, valine, phenylalanine, and tyrosine, indicating a family-wide lack of complete biosynthetic pathways for these amino acids. This information could be used to generate strain-specific minimal media and screen for wild-type auxotroph strains as a predictive tool to prevent the backslopping of industrially important strains.

FBA analysis was also performed to screen for the biotechnological potential of the species in PanGEM. As expected, PanGEM showed high production rates for lactate and acetate, which are well-known as primary organic acids produced by Lactic acid bacteria (32). PanGEM predicted the production of 41 compounds across all strains, some of which are biotechnologically important compounds, consistent with reports in the literature. These include acetoin (33), acetaldehyde (34), pyruvate (35), succinate (36), D-alanine (37), mannitol (38), formate (39), malate, citrate (39), propanediol (40), butanediol (41) and ethanol (42), as well as the neurotransmitter GABA (43), and vitamins such as riboflavin (44) and pyridoxal phosphate (45).

Using PanGEM, we investigated the biotechnological capabilities of product-specific isolates and their link to the organoleptic properties of the final product. GEMs for Kimchi, Tibet Kefir, Gruyere, and red wine isolates were analyzed, and PanGEM predicted the fermentation profile of related strains to identify their role in the quality of the final product. PanGEM contained 52 Kimchi isolates belonging to ten different species predicted to be capable of producing 15 different metabolites, including succinate responsible for umami taste (46), acetic acid as the main organic acid (47), acetaldehyde (48), and acetoin (49). Moreover, PanGEM predicted GABA production, which is one of the targeted metabolites for overproduction during Kimchi fermentation(50). Also, red wine isolates, consisting of 15 *Oenococcus onei* strains, were predicted as producers of malate (51), pyridoxine (52), and ornithine (53,54). The Kefir isolates comprised 14 strains from various species, with FBA predicting the production of 13 metabolites, including ethanol (55), succinate (56), acetoin (57), and lactic acid (57). D-alanine production was high in Kefir isolates, potentially contributing to the sensory quality. Gruyere isolates all belonged to *L. paracasei*, predicted to be capable of the production of 11 different metabolites, including acetoin (58) and succinate (59). Interestingly, a high production rate for D-alanine was predicted, which could be related to the sweet taste of Gruyere (23). Six isolates out of 11 were D-alanine producers.

The economic impact of *Lactobacillaceae* is enormous, enabling industries with an annual turnover exceeding a trillion U.S. dollars. Our study presents a family-wide metabolic reconstruction of *Lactobacillaceae* using pangenome analysis and metabolic reconstruction, resulting in high-quality genome-scale metabolic models. The validated PanGEM enables the discovery and understanding of the biotechnological potential of these bacteria, developing novel applications and screening for disease-associated microbial metabolites in the food industry. PanGEM provides a structured and computable genetic basis for metabolic traits in the *Lactobacillaceae* family.

## Methods

### 1. reactome reconstruction

#### 1-1. Data collection

For reconstruction of the *Lactobacillaceae-*specific reactome, 49 reference genomes (Supplementary File.1) were downloaded from NCBI covering nine genera and 33 species to capture the metabolic diversity of *Lactobacillaceae* as much as possible.

#### 1-2. Genome re-annotation

Initially, all genomes were annotated using the Prokka software (60), and stringent parameters were applied (Seemann, 2014). A group of 13 high-quality manually annotated genomes was selected from the NCBI Genbank database as reference genomes for annotation (Supplementary File.4). The output genbank files were merged into a single Genbank file and used as the reference genome of the reactome.

#### 1-3. GPR formulation and curation

Metabolic functions were assigned to the reference genome of the reactome using the Modelseed(61) pipeline to formulate the first draft of reactome. Subsequently, All reactions went through a manual curation process. A Cross-validation was performed for each reaction against KEGG universal reactome. GPR boolean rules were checked against BioCyc(62) and KEGG databases (63). Reactions and metabolite identifiers were mapped against the BIGG database(64). The directionality of each reaction was checked based on a) Gibbs free energy retrieved from Biocyc, b) BIGG reactions due to the availability of information, c) previous *Lactobacillaceae* GEM reconstructions (14,19,65–68). Metabolite charges were curated based on a) BIGG metabolites, b) KEGG metabolites, c) ChEBI metabolites d) previous GEMs. Where information was inconsistent across databases, Marvine’s suite was deployed to calculate metabolite charges at physiological pH (7.2). Reactions mass and charge balance were checked using the Cobrapy package (69) and were curated where necessary. Cellular subsystems were assigned to the reactions based on KEGG subsystems.

#### 1-4 Reactome refinement

The curated reactome was subjected to an iterative refinement process; for this goal a) spontaneous reactions (non-enzymatic reactions) were extracted from KEGG, BIGG, and previous reconstructions and were added to the draft reactome. b) exchange reactions were extracted from BIGG and previous reconstructions and were added to the draft reactome. c) genes missing from the reactome were identified and assigned to their corresponding reactions manually to form a GPR and included in the draft reactome. d) a bidirectional blast with a similarity threshold of 60% was performed on the reference genome of reactome against BIGG reconstructions to find and add further missing reactions. e) a Biomass objective function (BOF) was taken from *L. plantarum* reconstruction (68) and was modified based on information extracted from literature to formulate a Universal *Lactobacilli*-BOF. f) refined reactome was converted to a mathematical model using Cobrapy. g) a primary gapfilling was performed on the reactome to find and fill metabolic gaps (Supplementary Figure 7) within the reactome using fastgapfill algorithm (70). The output of this step is called *Lactobacillaceae* reactome hereafter and was used as a template for strain-specific GEM reconstruction across *Lactobacillaceae* members.

### 2-Lacto PanGEM reconstruction

#### 2.1 Genomes collection and re-annotation

*Lactobacillaceae* genomes were downloaded from NCBI. following quality control steps were performed during genomes selection (6):

1. 4,783 genomes of the *Lactobacillaceae* family were retrieved from the NCBI database.
2. Genome Taxonomy Database Toolkit (GTDB-Tk) was deployed to re-annotate the taxonomy for all 4,783 genomes (71).
3. Further, quality control and quality assurance (QC/QA) were done to get good-quality genomes. The QC/QA includes the taxonomy, number of contigs (<200), and N50(>50,000).
4. Species with less than 30 genomes were excluded to maintain sample distribution.
5. Finally, a total number of 2446 high-quality genomes of the *Lactobacillaceae* family remained to be passed to the Multi-strain GEM reconstruction workflow as input.

#### 2-2 Multi-strain reconstruction

GPR mapping was conducted using bidirectional best hits (BBH) with a similarity threshold of 70%. For this goal, quality-controlled genomes from the previous step were used as the target genome, and the reference genome of reactome was used as a reference. The resultant homology matrix was transformed into GPR presence-absence matrix (72). Mapped GPRs for each strain were collected to generate draft GEMs. Subsequently, all orphan, exchange, and spontaneous reactions, along with the *lactobacilli*-specific biomass reaction, were added to all drafts.

#### 2.3 Gapfilling

Gapfilling analysis was performed on generated GEMs to ensure their functionalities. Reactome was used as a collection of candidate reactions for gapfilling; all GEMs were gapfilled on CDM. initially, GEMs were gapfilled using fastgapfill, although it couldn’t return a possible solution for the majority of GEMs. Subsequently, an alternative methodology (Supplementary Figure 7), enabling large-scale gapfilling in a feasible timespan, was developed and applied.

### 3-Lacto PanGEM Analysis

#### 3-1 Dependencies

All analyses were performed in Python language deployed on an Azure virtual machine, statistical analysis was done using Python packages, including pandas, numpy and scipy, and figures were generated using Matplotlib and Plotly. Cobrapy was used for GEM analysis. In all FBA-based simulations, biomass reaction was defined as an objective function.

#### 3-2 Niche classification

Isolation sources were collected from NCBI biosample when available, and a total number of 608 distinct isolation sources were identified and classified into 9 larger groups including, plant-based, meat-based, dairy-based, commercial, commensal, environmental, beverages, uncategorized (undefined) food, and not reported (Supplementary Figure 1).

#### 3-2 Defining family-wide and species-specific core, accessory, and rare reactome

A reaction presence-absence matrix was constructed based on the *Lactobacillaceae* reactome and the reaction content of each strain. Family-wide Core (F>99%), accessory (15≤F≤99), and rare (0<F<15) reactome were defined based on reaction frequency (F) across PanGEM. Similar parameters were considered for species-specific core, accessory, and rare reactome. Reactions that were exclusive to a species (whether they could be found in one or multiple strains across a species) were called unique reactions.

#### 3-3 Validation

Validation was carried out over three different data types, including carbon source utilization, auxotrophy, and growth rate.

#### 3-3-1 Carbon source utilization

Capability of three strains for growing on 21 different C-sources was predicted and compared to experimental results obtained from the literature. To simulate growing on different C-sources, the exchange reaction of Glucose was bound to zero, and the lower bound of the exchange reaction of the target C-source was bound to 1000 mmol/gDCW/h. Growth was reported if the predicted growth rate was above 0.01/h. FBA was used for growth stimulation.

#### 3-3-2 Auxotrophy

Auxotrophy prediction was performed over six strains by bounding the lower bound of each of the 49 CDM components to zero. Auxotrophy was reported for a component If the predicted growth rate was below 0.01/h. FBA was used for growth stimulation.

#### 3-3-3 Growth rate

Growth rate was predicted for five strains using FBA on CDM. predicted growth rates were compared to experimental results obtained from the literature.

#### 3-3-4 Prediction of fermentation profile

Fermentation profile for each strain within PanGEM was predicted by FBA on CDM. strains were grouped based on isolation sources and species. Strains that were isolated from four industrially important food products were chosen to predict their fermentation profile and find a link between their metabolic capability and the organoleptic properties of their isolation source. A set of Metabolites whose exchange reaction was above zero was reported as a fermentation profile for each strain.

### 4-Availability of Scripts and Data

All data and scripts available at github.com/omidard/LactoGEMs.

## Notes

https://github.com/omidard/LactoPanGEM/blob/916ff8d18623bfad6519d5ed48643516562f59ec/Supplementary_File_2_Lactobacilli_Reactome.json

### Competing Interest Statement

The authors have declared no competing interest.

