## Supplementary materials for "Pangenome reconstruction of *Lactobacillaceae* metabolism predicts species-specific metabolic traits"

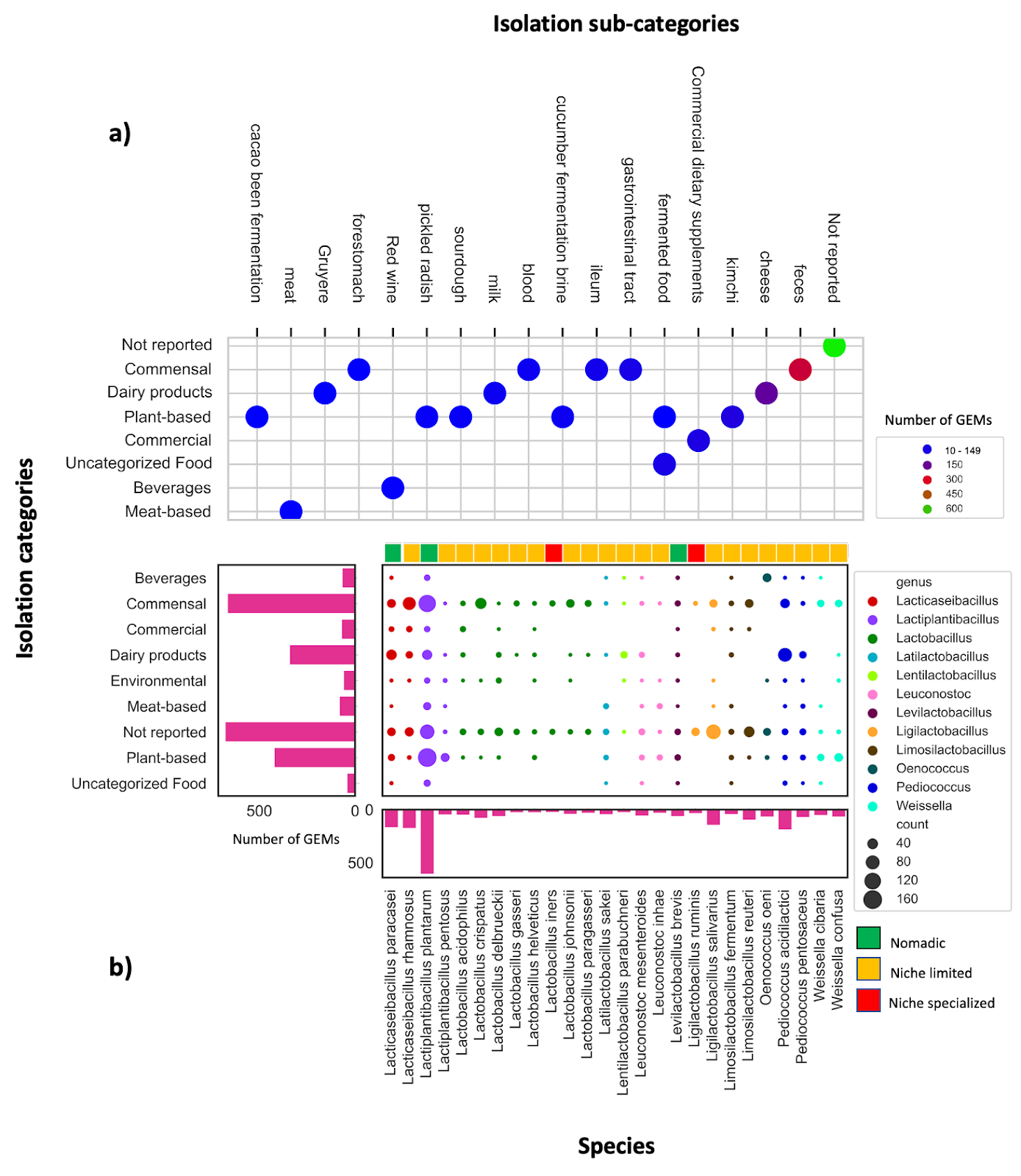

***Supplementary Figure 1: Scale and diversity of PanGEM for Lactobacillaceae****. Panel (a) displays the distribution of GEMs within sub-categories for each major isolation category, revealing significant diversity within the Lactobacillaceae. Panel (b) further highlights the diversity of 2,447 GEMs across nine different isolation sources, 26 species, and multiple genera. The y-axis shows major isolation sites, with colored dots representing GEMs belonging to each genus. The size of the dots corresponds to the number of isolates, while the color bar at the bottom indicates the niche specialization status of each species based on the number of unique isolation categories. The marginal plot on the x-axis shows the number of GEMs within each species, while the marginal plot on the y-axis depicts the number of GEMs within each major isolation source. This figure underscores the vast diversity of Lactobacillaceae and their potential for various applications in industry and biotechnology.*

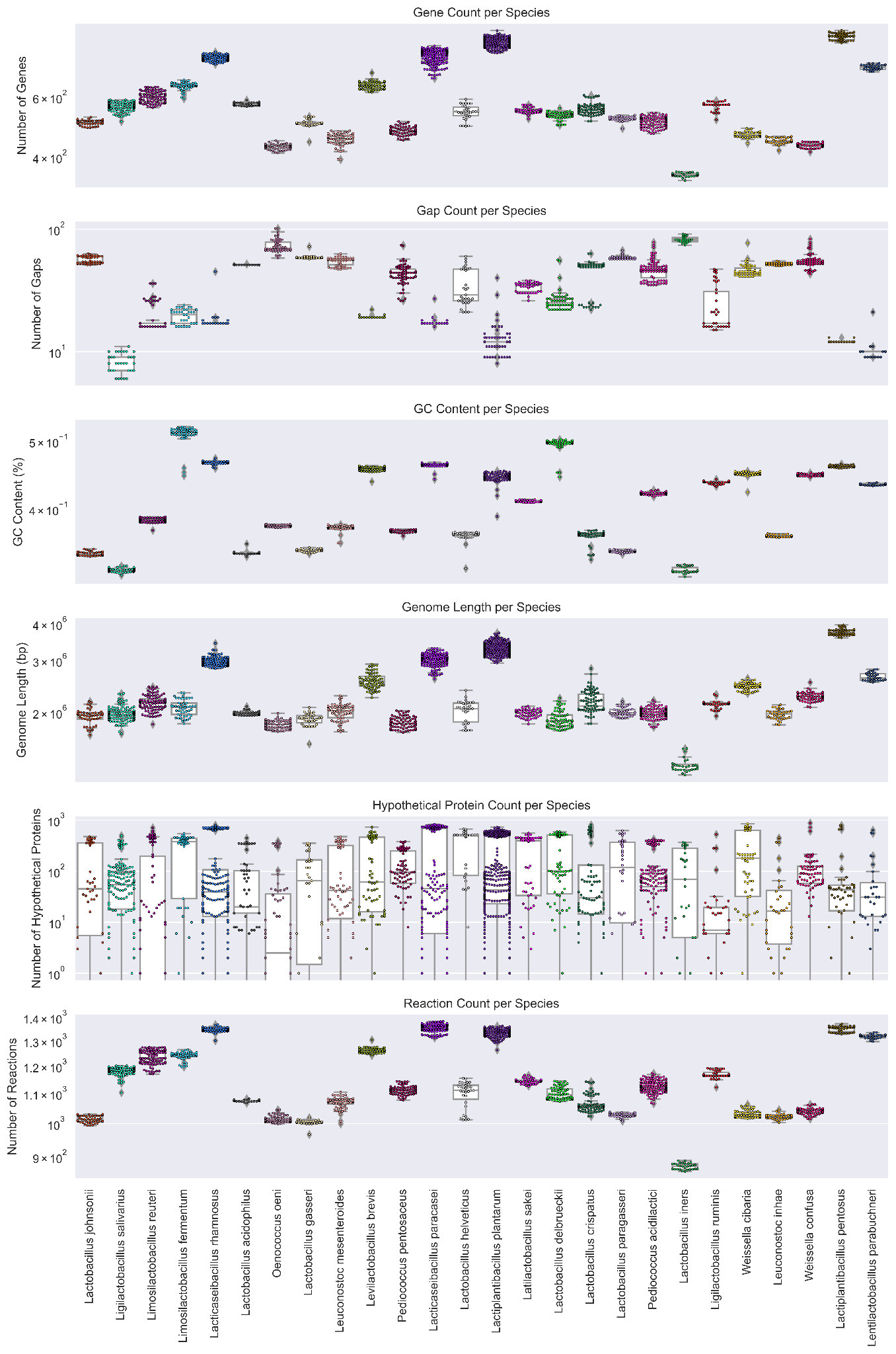

***Supplementary Figure 2: Comparative Analysis of six Genome and GEMs Characteristics Across Lactobacillaceae****. The figure consists of six subplots depicting the distribution of various genome characteristics across multiple bacterial species. The subplots include (A) reaction count, (B) gene count, (C) gap count, (D) GC content, (E) genome length, and (F) number of hypothetical proteins. Each boxplot shows the distribution of the respective GEMs characteristics across the selected bacterial species. The x-axis in each subplot shows the name of the bacterial species, while the y-axis represents the respective characteristic measured.*

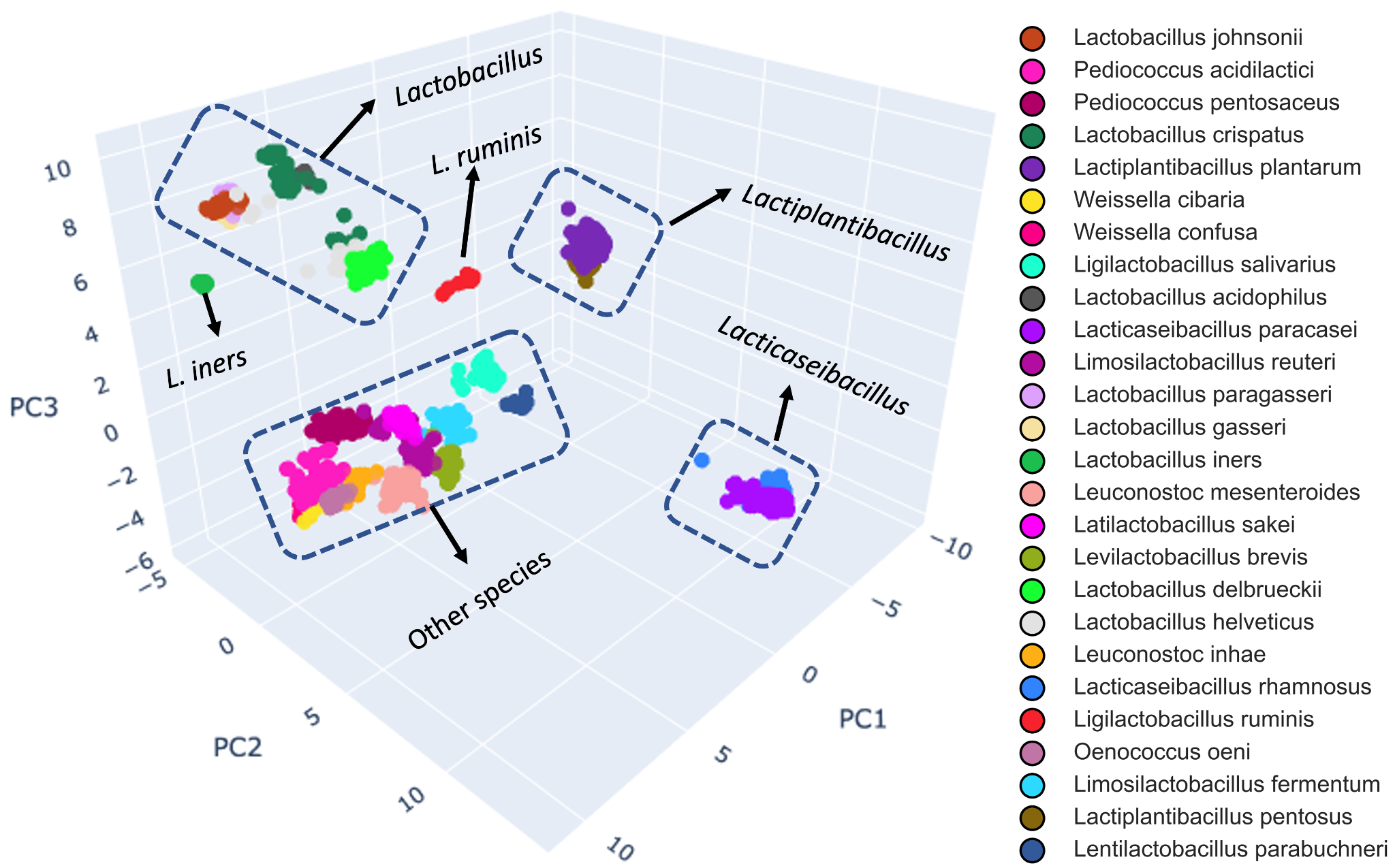

***Supplementary*** ***Figure 3:*** *Three-dimensional PCA plot of 2446 bacterial strains based on their metabolic reactions, color-coded by species. The plot reveals six distinct clusters (Supplementary Note 3), including one each of Lactobacillus, Lactoplantibacillus, and Lactocaseibacillus genus, as well as a separate cluster for Lactobacillus ruminis and Lactobacillus iners, and a final cluster containing all remaining species.*

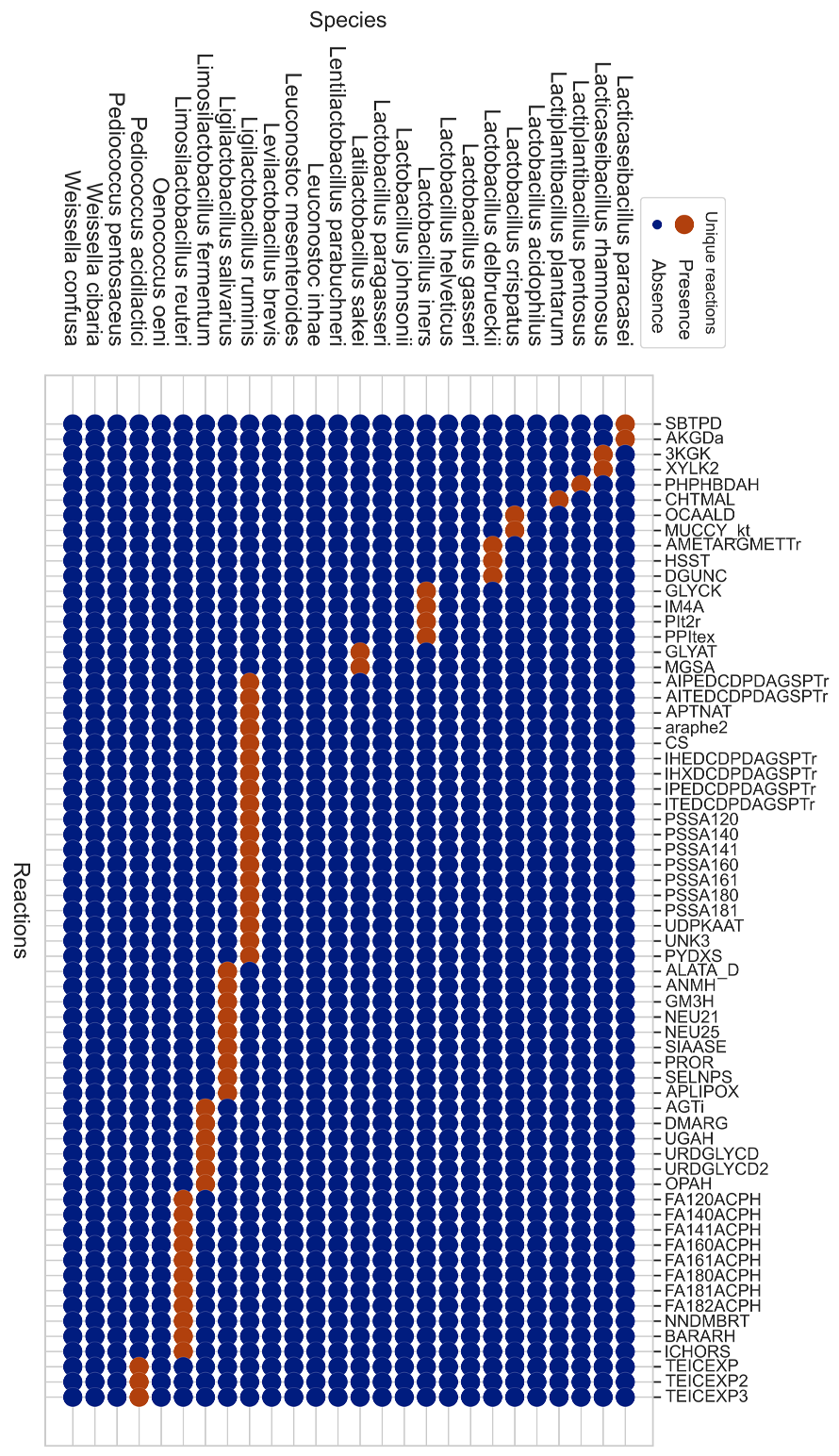

***Supplementary Figure 4: Lactobacillaceae unique reactions.*** *The scatter plot shows the distribution of unique reactions across 26 species of Lactobacillaceae PanGEM, with species-specific (unique) reactions represented by brown dots. L. ruminis stands out as having the highest number of unique reactions (19) among the species examined, while L. reuteri, L. salivarius, and L. fermentum follow with the next highest number of unique reactions. The scatter plot highlights the unique metabolic capabilities of Lactobacillaceae. Lactobacillus reuteri, Lactobacillus salivarius, and Lactobacillus fermentum are commonly found in the human gastrointestinal tract and oral cavity, as well as in other mammalian species. This would suggest that the acquisition of new metabolic capabilities may be an important mechanism for bacteria to adapt and persist in complex environments, such as the human gut. Therefore, the high number of unique reactions observed in these Lactobacillus species may be indicative of their ability to acquire and integrate new metabolic functions from other members of the gut microbiota.*

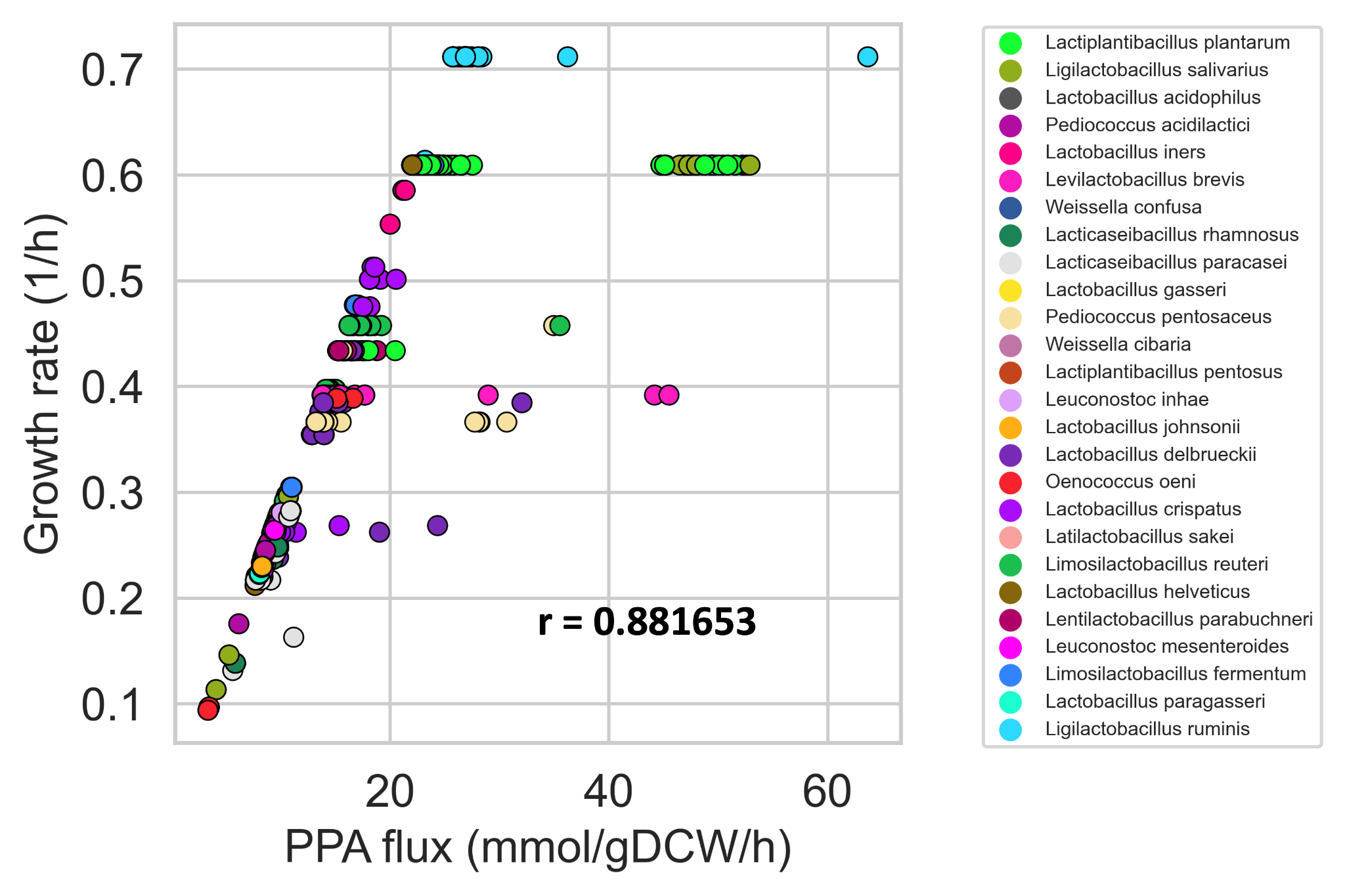

***Supplementary Figure 5: Positive Correlation Between PPA Enzyme Flux and Growth Rate Across Lactobacillaceae PanGEM Species.*** *the scatter plot shows a strong positive correlation (r=0.88) between PPA enzyme flux and growth rate across multiple species. Each data point represents a distinct species, color-coded for clarity. The results suggest that an increase in PPA enzyme flux is associated with a higher growth rate, highlighting the importance of PPA enzyme regulation in cellular metabolism.*

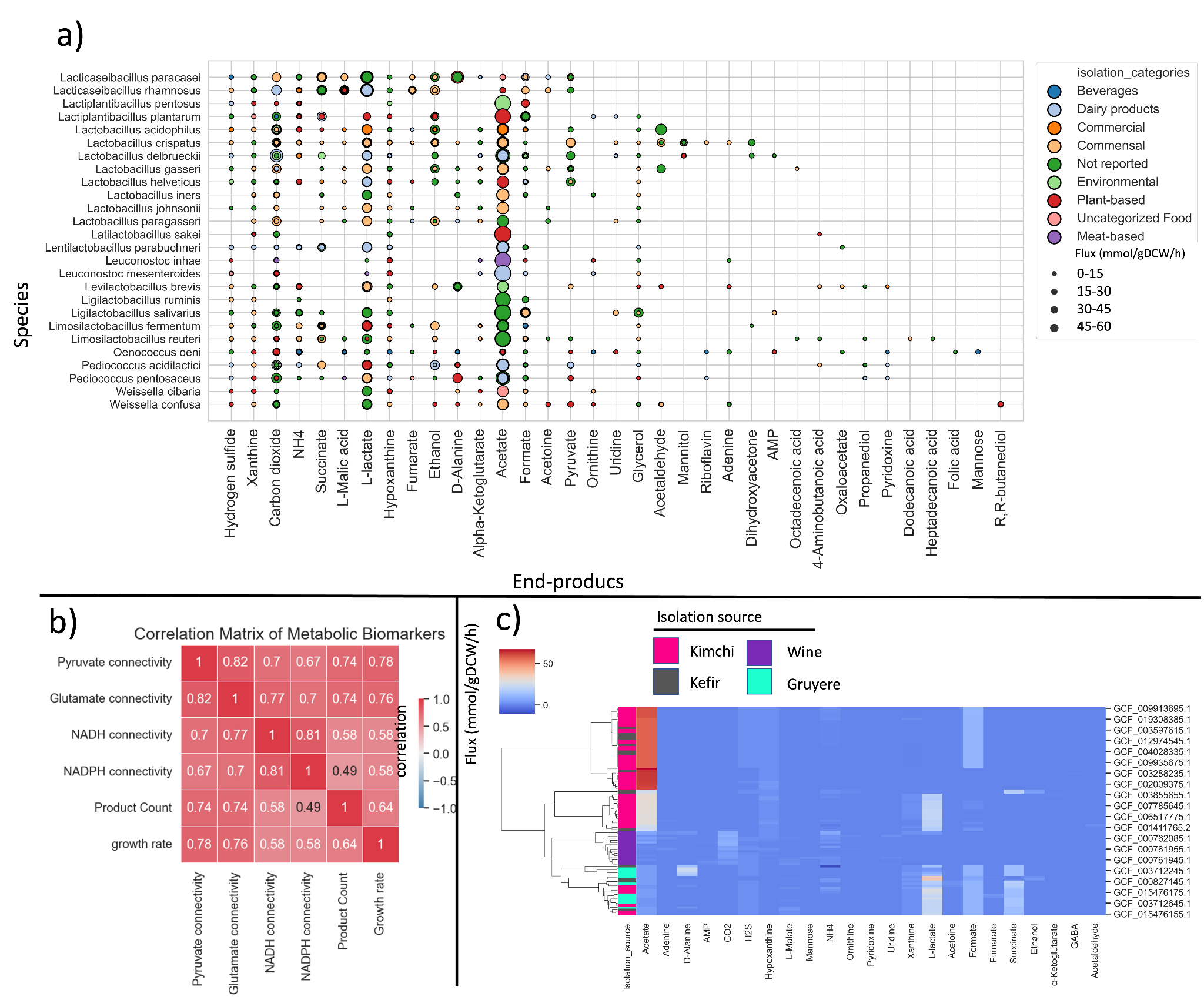

***Supplementary Figure 6: by-product formation prediction****.* ***a)*** *by-product formation across Lactobacillaceae dispersed over isolation categories. Isolation categories are color coded and dot size encodes the production rate of by-products (mmol/gDCW/h).* ***b)*** *Heatmap shows correlations between NADH connectivity, NADPH connectivity, Pyruvate connectivity, Glutamate connectivity, growth rate, and number of by-products across 2446 Lactobacillaceae strains. Pyruvate and Glutamate connectivity show a strong positive correlation (r=0.74) with the number of by-products produced, highlighting the importance of these metabolites in determining the metabolic diversity of Lactobacillaceae.* ***c)*** *Cluster map depicts product formation rate predicted by Lactobacillaceae PanGEM for kimchi, gruyere, and red wine isolates. The majority of strains show clustering based on their isolation source, indicating that these strains are responsible for different organoleptic features of the by-products. Notably, strains isolated from kimchi exhibit a high production rate of acetate, suggesting that the metabolic activity of these strains contributes to the distinctive flavor and aroma of kimchi. These findings highlight the importance of strain selection in the production of fermented foods with desirable organoleptic properties.*

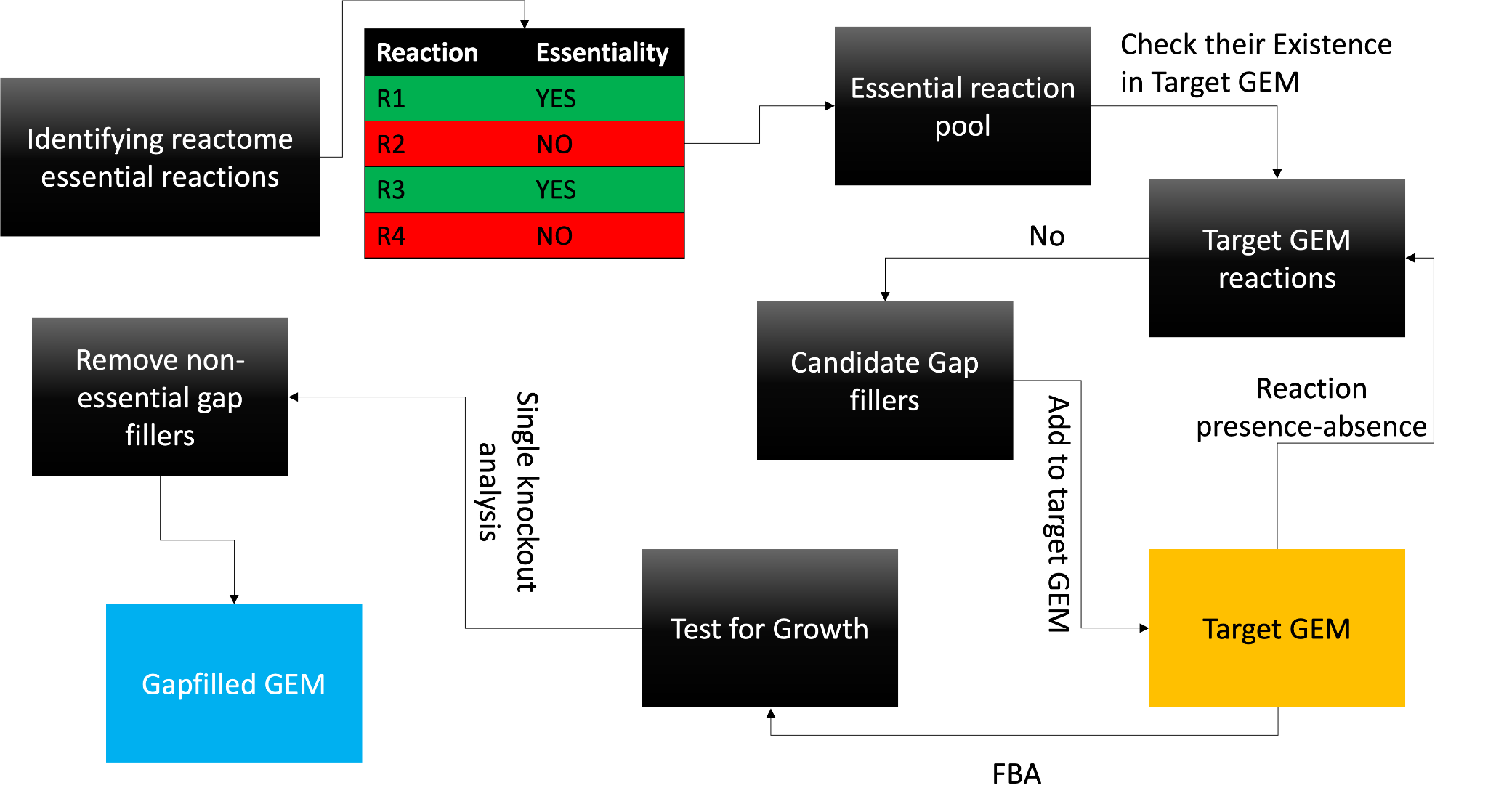

***Supplementary Figure 7: Schematic diagram of the gapfilling procedures used for the reconstruction of 2446 GEMs****. The gapfilling process involved identifying the essential reactions in a reactome and constructing an Essential Reaction Pool (ERP). The target GEM was then checked for the presence of these essential reactions. If any essential reactions were missing, they were added to the target GEM. After gapfilling, growth simulations were performed, and non-essential gapfillers were removed using single knockout simulations. This gapfilling approach enabled the reconstruction of comprehensive and accurate GEMs for a large number of Lactobacillaceae strains, facilitating in-depth metabolic analysis and optimization of these strains for industrial applications.*

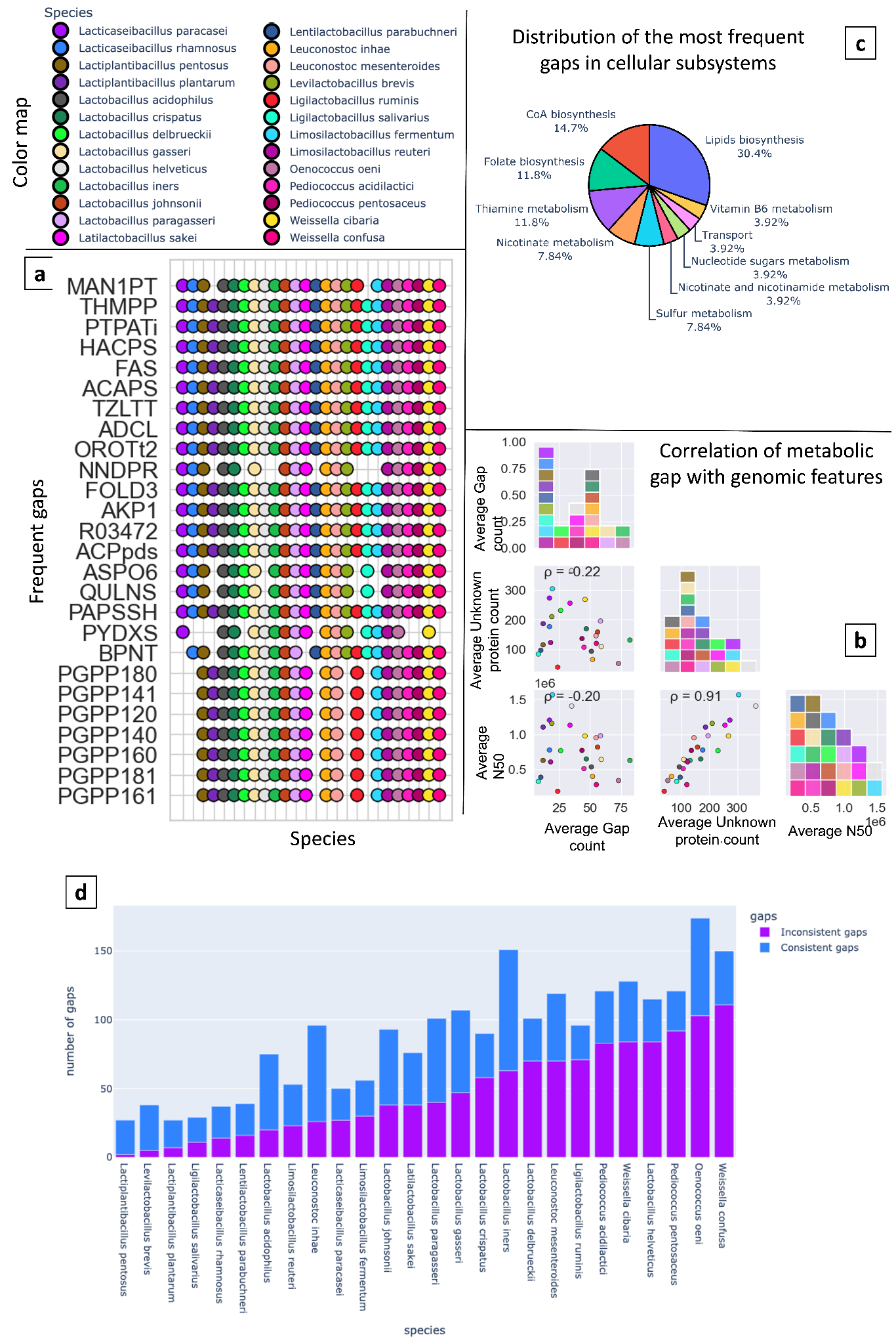

***Supplementary Figure 8: Gap analysis and gap frequency within Lactobacillaceae PanGEM****.* ***a)*** *Most common metabolic gaps within Lactobacillaceae PanGEM, categorized based on gaps frequency in each species. Species are shown on Y-axis, and X-axis shows reactions that have been identified as gaps and have been added to GEMs. Colored dots show the presence of each gap within GEMs, and each color is assigned to a species (color map). b) correlation of gap count with genome contiguity (N50) and knowledge/technical gap (Unknown protein). Colors are coded based on the color map. c) distribution of the most common gaps within cellular subsystems. Values are presented as percentages. d) Stacked bar chart showing consistent and inconsistent gaps in Lactobacillaceae pan-genome models (Lactobacillaceae PanGEM). The y-axis represents the gap count, and the x-axis shows the species analyzed. Inconsistent gaps are present in some, while consistent gaps are present in all strains of a species*

***Table. 1. Exchange reaction constraints for CDM simulation by FBA******

| *Exchange Reaction* | *Lower Bound (mmol/gDCW/h)* | *Compound Name* |
| --- | --- | --- |
| *Carbon source* | | |
| *EX_glc_D_e* | *-25,2* | *D-Glucose* |
| *EX_ac_e* | *-1* | *Acetate* |
| *EX_cit_e* | *-1* | *Citrate* |
| *Amino acids* | | |
| *EX_arg_L_e* | *-1* | *L-Arginine* |
| *EX_cys_L_e* | *-1* | *L-Cysteine* |
| *EX_glu_L_e* | *-2* | *L-Glutamate* |
| *EX_ile_L_e* | *-1* | *L-Isoleucine* |
| *EX_leu_L_e* | *-1* | *L-Leucine* |
| *EX_met_L_e* | *-1* | *L-Methionine* |
| *EX_tyr_L_e* | *-1* | *L-Tyrosine* |
| *EX_phe_L_e* | *-1* | *L-Phenylalanine* |
| *EX_thr_L_e* | *-1* | *L-Threonine* |
| *EX_val_L_e* | *-1* | *L-Valine* |
| *EX_gly_e* | *-1* | *Glycine* |
| *EX_ala_L_e* | *-1* | *L-Alanine* |
| *EX_asp_L_e* | *-2* | *L-Aspartate* |
| *EX_his_L_e* | *-1* | *L-Histidine* |
| *EX_lys_L_e* | *-1* | *L-Lysine* |
| *EX_pro_L_e* | *-1* | *L-Proline* |
| *EX_ser_L_e* | *-1* | *L-Serine* |
| *EX_trp_L_e* | *-1* | *L-Tryptophan* |
| *Nucleotides* | | |
| *EX_ura_e* | *-1* | *Uridine* |
| *EX_gua_e* | *-1* | *Guanosine* |
| *EX_ins_e* | *-1* | *Inosine* |
| *EX_ade_e* | *-1* | *Adenosine* |
| *EX_xan_e* | *-1* | *Xanthine* |
| *EX_orot_e* | *-1* | *Orotate* |
| *Vitamines* | | |
| *EX_btn_e* | *-1* | *Biotin* |
| *EX_pnto_R_e* | *-1* | *Pantothenate* |
| *EX_thm_e* | *-1* | *Thiamine* |
| *EX_pydam_e* | *-1* | *Pyridoxamine* |
| *EX_pydxn_e* | *-1* | *Pyridoxine* |
| *EX_ribflv_e* | *-1* | *Riboflavin* |
| *EX_fol_e* | *-1* | *Folate* |
| *EX_ascb_L_e* | *-1* | *L-Ascorbate* |
| *EX_4abz_e* | *-1* | *4-Aminobenzoate* |
| *EX_nac_e* | *-1* | *N-Acetyl-D-glucosamine* |
| *EX_thymd_e* | *-1* | *Thymidine* |

** The constraints applied to the exchange reactions for Flux Balance Analysis were kept consistent throughout the entire study*

**Table. 2. Global market size of *Lactobacillaceae-related* products**

| **Product** | **Market size (billions of U.S. dollars)** | **Reference** |
| --- | --- | --- |
| **Yogurt** | **167** | **(Yogurt - worldwide )** |
| **Cheese** | **154.8** | **(Baron )** |
| **Butter** | **51.49** | **(Straits Research )** |
| **Sour cream** | **1.7** | **(Sour Cream Market 2023 )** |
| **Kefir** | **1.3** | **(Kefir Market )** |
| **Soy sauce** | **40.63** | **(Soy sauce market size, share, trends )** |
| **Red wine** | **182.0** | **(Red Wine Market )** |
| **Miso** | **107.4** | **(Miso market )** |
| **Tempeh** | **4.7** | **(Tempeh Market )** |
| **Kimchi** | **3.3** | **(Kimchi Market size, report, statistic...)** |
| **Sauerkrauts** | **10.4** | **(Sauerkrauts Market size, share, trend...)** |
| **Pickles** | **11.1** | **(Research and Markets ltd )** |
| **Kombucha** | **2.64** | **(Kombucha market size, share & trends ...)** |
| **Sourdough** | **3.13** | **(Sourdough Market - industry analysis ...)** |
| **Vinegar** | **6.4** | **(Vinegar market size, share & trends a...)** |
| **Pepperoni** | **2** | **(Pepperoni market )** |
| **Salami** | **7.5** | **(Reshovski 2022)** |
| **Fermented fish** | **66.9** | **(Processed fish market )** |
| **Chocolate** | **113.16** | **(Chocolate market size, share & trends...)** |
| **Probiotic Supplements** | **6.65** | **(Global Probiotic Dietary Supplements ...)** |
| **Whey protein powder** | **12.41** | **(Precedence Research 2023)** |
| **Lactic acid** | **3.1** | **(Lactic acid market size, share & tren...)** |
| **Hot Sauce** | **2.75** | **(Facts and Factors 2022)** |
| **Feed probiotics** | **4.8** | **(MarketsandMarkets Research Pvt. Ltd. ...)** |
| **Total market size** | **938** |  |

**Supplementary Note 1**

A species-specific reactome analysis was performed to understand metabolic conserveness among strains of each species. For this goal, core, accessory, and rare reactomes were calculated for each species based on intra-species reactions commonality. The highest percentage of rare reactome was found for *L. crispatus, L. mesentroides, O. onei,* and *L. fermentum.* On the contrary, *L. iners, L. ruminis*, and *L. acidophilus* had the lowest percentage of rare reactome. Also, the highest percentage of accessory reactome could be found in *L. plantarum, L. helveticus,* and *P. acidilactici. In contrast,* the lowest percentage of accessory could be found in *L. acidophilus, L. iners,* and *L. parabuchneri.* Core reactome showed a high percentage in *L. acidophilus, L. iners, L. parabuchneri,* and *L. ruminis,* while *L. plantarum, L. helveticus, L.crispatus and P.acidilactici.* This shows *L. plantarum* has the most metabolically diverse strains across the whole family, while *L. acidophillus* is the most metabolically conserved species.

**Supplementary Note 2**

To reveal additional metabolic similarities and differences of *Lactobacillaceae*, the essentiality of CDM media components for each GEM has been predicted by FBA (Fig.3-e). This analysis showed that *Lactiplantibacillus plantarum,* with nine consistent auxotrophies among all its members and one inconsistent auxotrophy (Cysteine predicted to be essential in some of *Lactiplantibacillus plantarum* but not all), has the lowest number of auxotophies among all *Lactobacillaceae*, despite *Pediococcus acidilactici* and *cibaria* which have the highest number of auxotrophies, 12 consistent auxotrophies were predicted for both species, while cibaria has eight more inconsistent auxotrophies and this number for *acidilactici* predicted to be seven. Also, regarding the highest number of consistent auxotrophies, *Latilactobacillus sakei,* and *Lactobacillus paragasseri,* with 16 consistent auxotrophies among all strains within these two species, have the highest number of consistent auxotrophies (Fig3.e). Also, four amino acids, including Isoleucine, Valine, Phenylalanine, and Tyrosine, were predicted as globally essential for all 2,446 models in *Lactobacillaceae* PanGEM.

Supplementary Note 3

Distinct metabolic profiles were observed among the different *Lactobacillaceae* species based on the clustering patterns seen in the PCA plot. These metabolic differences likely indicate adaptations to various ecological niches as the bacteria must obtain the necessary nutrients for survival and growth in their environment. Differences in preferred ecological niches were apparent in the distinct clustering of *Lactobacillus, Lactiplantibacillus*, and *Lacticaseibacillus* species. *Lactobacillus* species are commonly found in the gastrointestinal tracts of humans and animals. In contrast, *Lactiplantibacillus* species thrive in plant environments such as soil, plant surfaces, and fermenting vegetables. *Lacticaseibacillus* species are known to prefer dairy environments, such as milk and cheese. Notably, a separate cluster of *Lactobacillus ruminis* was observed, suggesting adaptation to a specialized ecological niche within the digestive tracts of cattle and other ruminants. The observed clustering patterns suggested a close connection between bacterial species' metabolic diversity and ecological niches. Further understanding of these metabolic differences and ecological niches can provide valuable insights into the functional roles of these bacteria in different environments and their potential biotechnological applications.
